# Elp1 is required for development of visceral sensory peripheral and central circuitry

**DOI:** 10.1101/2022.04.11.487663

**Authors:** Zariah Tolman, Marta Chaverra, Lynn George, Frances Lefcort

## Abstract

Cardiovascular instability and a blunted respiratory drive in hypoxic conditions, are hallmark features of the genetic sensory and autonomic neuropathy, familial dysautonomia (FD). FD results from a mutation in the gene *ELP1*, whose encoded protein is a scaffolding subunit of the six subunit Elongator complex. In mice, we and others have shown that *Elp1* is essential for the normal development of neural crest derived-dorsal root ganglia (DRG) sensory neurons. Whether *Elp1* is also required for development of ectodermal placode-derived visceral sensory receptors which are required for normal baroreception and chemosensory responses, has not been investigated. Using mouse models for FD, our data indicate that in fact the entire circuitry underlying baroreception and chemoreception is impaired due to a requirement for *Elp1* not only in the visceral sensory neuron ganglia, but also for normal peripheral target innervation, and in their CNS synaptic partners in the medulla. Thus *Elp1* is required in both placode- and neural crest-derived sensory neurons and its reduction aborts the normal development of neuronal circuitry essential for autonomic homeostasis and interoception.

**Summary statement:** Due to faulty afferent sensory signaling, patients with Familial dysautonomia (FD) have a diminished sensory arm of the baroreflex which would normally modulate blood pressure, and they have a blunted response to hypoxia and hypercapnia (Norcliffe-Kaufmann et al. 2010). Using mouse models for FD, we reveal here the underlying pathology which may underlie these severely impaired homeostatic reflex pathways in FD.

## Introduction

Familial dysautonomia (FD), also referred to as Hereditary sensory and autonomic neuropathy Type III, is a devastating rare genetic neuropathy that results from mutations in the gene, *ELP1* (previously referred to as *IKBKAP*). Over 99.5% of patients inherit the identical T-C founder mutation that weakens inclusion of exon 20, causing a frame shift, and nonsense-mediated decay of the truncated mRNA (Slaugenhaupt et al. 2001; Cuajungco et al. 2003; Ibrahim et al. 2007). The dysfunctional splicing is cell type specific, with neurons being least capable of splicing the mutated pre-mRNA and hence being the cell type most impaired in FD (Slaugenhaupt et al. 2001; Cuajungco et al. 2003). The Elp1 protein is a key scaffolding subunit of the six-subunit Elongator complex which is essential for tRNA wobble uridine (U34) modifications; in the absence of a functional elongator complex, additions of thiol and methoxycarbonyl-methyl do not occur, which perturbs the translation of mRNAs that preferentially use either AA- or AG-ending codons (Huang et al. 2005; Bauer and Hermand 2012; Karlsborn et al. 2014; Kojic et al. 2021). Consequently, loss of Elp1 function leads to considerable changes in the cellular proteome, which can also indirectly perturb the cellular transcriptome, both of which negatively impact pathways that are critical for neurogenesis and survival (George et al. 2013; Chaverra et al. 2017; Ohlen et al. 2017; Goffena et al. 2018; Kojic et al. 2021).

While both the PNS and CNS are affected in FD, the major impaired cell type in this disease are sensory neurons, and in particular, visceral sensory neurons. For example, one of the most debilitating clinical hallmarks suffered by FD patients is cardiovascular instability which results from a faulty baroreflex and chemoreflex. These reflexes are vital for rapidly adjusting blood pressure, heart rate and respiration to maintain homeostasis. When blood pressure drops in healthy individuals, heart rate increases to increase blood pressure; in contrast, when blood pressure elevates, heart rate decreases to restore normal blood pressure levels. This reflex circuit is initiated by changes in activity of visceral sensory afferents whose cell bodies reside in the petrosal (cranial nerve IX) and nodose/jugular ganglia (cranial nerve X) (Kumada et al. 1990; Chapleau 2003; Benarroch 2008; Wehrwein and Joyner 2013; Min et al. 2019). The axons of the petrosal baroreceptors innervate the carotid sinus while the aorta is innervated by axons extending from nodose ganglion neurons (CNX). These same primary visceral sensory neurons project axons onto second order neurons in the nucleus of the solitary tract (NTS) in the medulla, the first central relay for visceral information (Blessing 1997; Kim et al. 2020). Dorsal to the NTS and also projecting onto it, is the area postrema (AP), a circumventricular organ whose neurons are in direct contact with the bloodstream and cerebrospinal fluid. In summary, visceral sensory information concerning blood pressure and levels of oxygen and carbon dioxide, are sent into the brainstem via cranial nerve IX and X visceral peripheral afferents, to activate compensatory changes in autonomic output, either through the vagus nerve or sympathetic nerves, to maintain homeostatic heart rate and blood pressure. This circuitry is illustrated in Figure 1.

**Figure 1.**
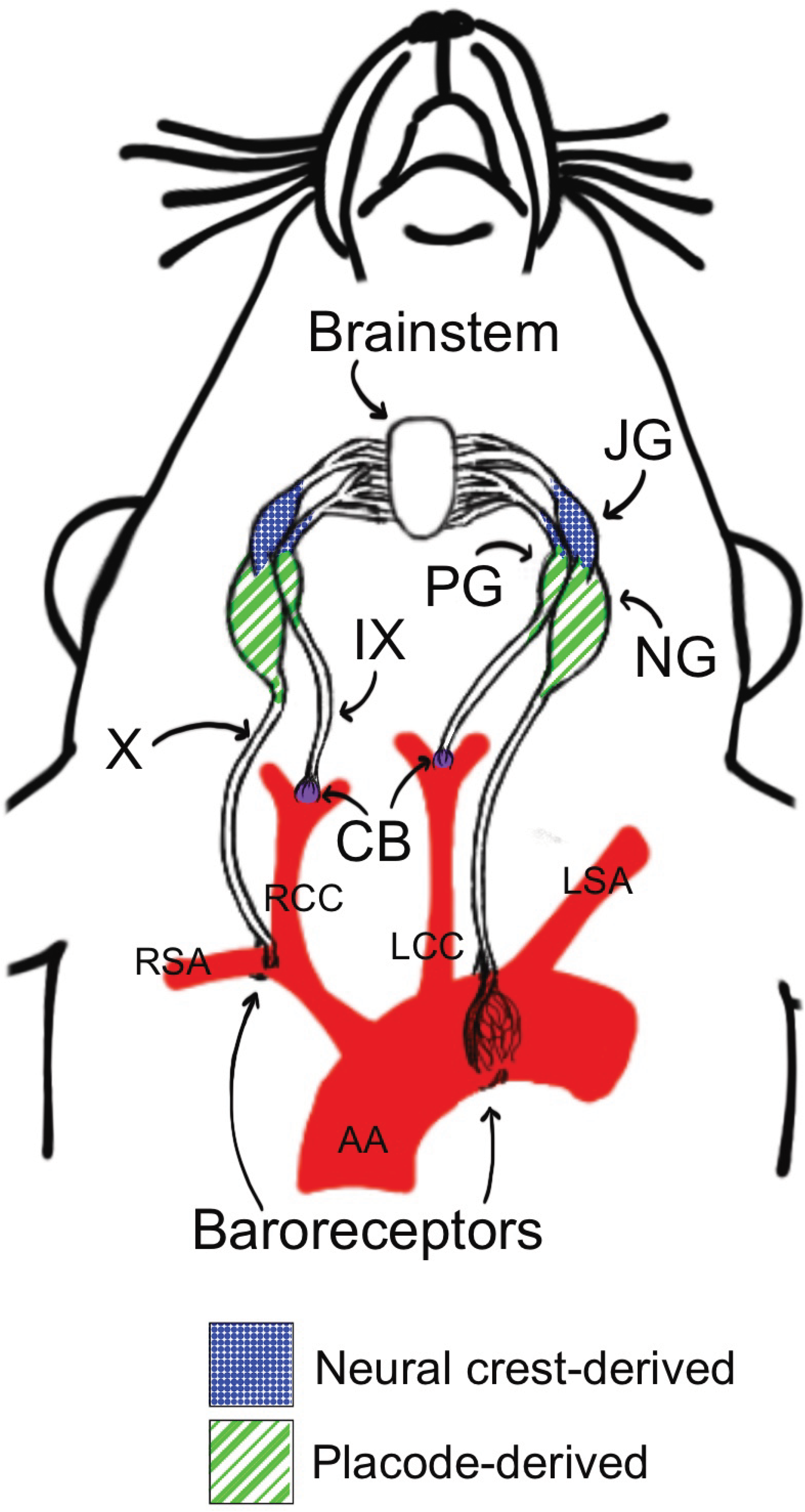
Cranial nerve IX and X circuits that mediate chemoreception and baroreception. The schematic depicts vagal sensory afferents that sense blood pressure through the aortic depressor nerve branch of the vagus nerve (X). The Glomus cells in the carotid body (purple, CB) relay chemosensory information concerning levels of O2, CO2, pH in addition to glucose, lactate and more, through the glossopharyngeal nerve (IX) to the brainstem. Baroreceptors sensing blood pressure in the aortic arch (AA) and right subclavian artery (RSA) relay this information to the nucleus of the solitary tract in the brainstem. The jugular ganglion (JG) of the vagus is neural crest-derived while the nodose ganglion (NG) and petrosal ganglion (PG) are ectodermal placode derived. These pathways are disrupted in FD. Other Abbreviations: RCC: right common carotid artery, LCC: left common carotid artery, LSA: left subclavian artery.

In addition to the mechanosensory baroreceptors that innervate the aorta, a second key visceral sensory population are chemoreceptors, which ensure homeostatic maintenance of ventilatory drive for respiration by detecting partial pressures of oxygen, CO2, and pH (Kameda et al. 2002; Ortega-Sáenz and López-Barneo 2020) for references). These chemoreceptive cells are located both peripherally in the carotid body, and centrally in the brainstem, in the Retrotrapezoid nucleus (RTN; Souza et al., 2019). The carotid body sits in the bifurcation of the carotid artery (Figure 1) and is composed of the sensory glomus cells (in addition to the supporting sustentacular cells), which express tyrosine hydroxylase and serotonin, and are derived from neuronal precursors in the adjoining sympathetic cervical ganglion (Kameda 2005). The glomus cells are multimodal sensory cells and relay essential information concerning blood oxygen, CO2, acid and glucose levels to the petrosal axonal afferents (Gonzalez et al. 1994; Pardal and López-Barneo 2002; Dauger et al. 2003). Through the glossopharyngeal nerve these afferents synapse in the nucleus tract solitarius (NTS) in the brainstem, and from there projections are sent to other brainstem nuclei that drive respiration, including the pre-Botzinger complex (Finley and Katz 1992).

Elegant studies in FD patients have demonstrated that the major point source of failure in their baroreflex is reduced sensory afferent signaling. Moreover FD patients have a blunted hypoxic ventilatory drive and do not respond normally to low levels of CO2 (Norcliffe-Kaufmann et al. 2010; Palma et al. 2019; Kaufmann et al. 2020). Both of these deficits underlie the frequent hypotension experienced by FD patients. Furthermore, FD patients suffer from swallowing abnormalities, which triggers aspiration and subsequent lung infections. In addition, they suffer from gastrointestinal, kidney and liver dysfunction, all tissues that are heavily innervated by the vagus nerve. In summary, the most life-threatening symptoms FD patients experience result from severe disruption in vagal and glossopharyngeal cranial nerve function.

The vagus nerve is the major visceral sensory and motor (parasympathetic) nerve, innervating all visceral organs to convey key physiological changes in the body to the CNS. The sensory neuronal cell bodies for this nerve are all located in cranial ganglia X, which include the nodose and jugular ganglia, which in the mouse tend to be fused with the petrosal ganglion (cranial ganglion IX) to form the nodose-petrosal-jugular complex (NPJ). This ganglia-complex in mammals derives embryologically from both neural crest and the epibranchial placodes (Harlow et al. 2011; Karpinski et al. 2016). Given the numerous vital functions subserved by vagal sensory neurons and the diversity of targets surveilled by them, several recent studies have transcriptionally profiled nodose/petrosal/jugular neurons and cells in the carotid body by single cell RNA sequencing (Peeters et al. 2006; Kupari et al. 2019; Mazzone et al. 2020; Prescott et al. 2020) to generate an atlas and map of the diversity of cell populations within the NPJ. Depending on the study, 18-37 classes of molecularly distinct nodose/petrosal/jugular population subsets were identified speaking to the considerable heterogeneity and diversity of this population. Neuronal subsets have also been identified and distinguished based on site of target innervation, function, and axonal terminal morpholog (e.g. (Chang et al. 2015; Williams et al. 2016; Hennel et al. 2018; Bai et al. 2019; Min et al. 2019; Prescott et al. 2020).

Given the profound role that the vagus nerve exerts in the body, and our incomplete understanding of which neuronal cell populations require Elp1, the goal of this study was to determine the fate of key visceral sensory neurons in mouse models for FD to better understand the pathophysiological mechanisms underlying the faulty baroreflex and chemoreflex of FD patients. This impairment could result from reduced neuronal number in the IX and X cranial ganglia, diminished target innervation by sensory axons, and/or reduced neuronal number in the key brainstem nuclei that receive and process visceral afferent input. Our study reveals that in fact, Elp1 is required in the entire circuit: in both the sensory and brainstem neurons, and is required for target innervation.

## MATERIALS AND METHODS

### Mice

All experiments with animals were performed according to the National Institutes of Health Guide for Care and Use of Laboratory Animals and were approved by the Montana State University Institutional Animal Care and Use Committee. The *Rosa^mTmG^ (Jackson Labs, Gt(ROSA)26Sor^tm4(ACTB-tdTomato,-EGFP)Luo^/J;* Strain #007576); *Elp1^LoxP/LoxP^, Tuba1a-Cre* (kind gift of Dr. Lino Tessarollo, NCI, Frederick, MD (Coppola et al. 2004)*); Elp1^LoxP/LoxP^, Wnt1-Cre (Jackson Labs, H2az2^Tg(^Wnt1-cre^)11Rth^ Tg(Wnt1-GAL4)11Rth/J;* Strain #003829*); Elp1^LoxP/LoxP^, and Elp1^BetaGal^ (Ikbkap^tm1a(KOMP)Wtsi^*, International Knock-out Mouse Consortium) mice have all been previously described by our lab (George et al. 2013; Chaverra et al. 2017). *Phox2b-Cre; Elp1^+/LoxP^* were generated in our lab using a *Phox2b-cre* line purchased from Jackson Labs (B6(Cg)-Tg(Phox2b-cre)3Jke/J) and crossed to our *Elp1^LoxP/LoxP^* mice to generate *Phox2b-cre; Elp1^LoxP/LoxP^* mice that occur at a 1 in 4 ratio from this cross, and have Elp1 deleted from Phox2b+ cells. All genotyping was performed via PCR.

#### LacZ staining

Mice that contained a *LacZ Ikbkap* reporter were obtained from the International Knockout Mouse Consortium *(Ikbkap^tm1a(KOMP)Wtsi^).* This allele contains an FRT-flanked beta-gal reporter cassette (lacZ) that disrupts the expression of *Elp1* but heterozygotes are used; these mice have no mutant phenotype (George et al., 2013). Tissue specific expression of the *Elp1:LacZ* reporter can be visualized by beta-galactosidase staining. Embryos were fixed (1% formaldehyde, 0.2% glutaraldehyde, and 0.02% NP-40 in PBS) for 30 minutes at room temperature. Standard LacZ staining was performed at 37°C overnight. Embryos were then cryoprotected in 30% sucrose/PBS and embedded in optimal cutting temperature compound (OCT; Sakura Finetek, Torrance, CA) for cryostat sectioning. Sections were analyzed with a light microscope.

### Immunohistochemistry

#### Aortic Arch Whole Mount Immunolabeling

Upper bodies of E18.5 or P1 mice were fixed in 4% paraformaldehyde (PFA) at 4° Celsius for 2.5 hours. Following 3 rinses in 1xPBS, aortic arches and surrounding cardiovascular tissue, including the bifurcation of the brachiocephalic artery into the right subclavian artery and right common carotid artery, were dissected. Tissue was blocked overnight in 1x animal-free blocker (AFB; Vector Laboratories, #SP-5030) with 0.5% Tween-20, 10% DMSO, and 2% glycine, followed by incubation in primary antibody for 3 nights. Tissue was then washed 3 x 15-minutes in block followed by overnight incubation in secondary antibody and 3 x 15-minute washes in 1xPBS. Tissue was then mounted in Vectashield or Prolong Gold.

Embryos were fixed in 4% paraformaldehyde for 2 hours at 4°C. After PBS washes, embryos were cryoprotected in 30% sucrose overnight at 4°C and embedded in OCT. Tissue was then sectioned at 14 um. For immunohistochemistry (IHC), sections were blocked AFB and 0.5% Triton X-100 for 1 hour at room temperature, then primary antibodies were applied and incubated at 4°C overnight. Primary antibodies used included tyrosine hydroxylase (TH) rabbit (1:1000, Millipore, #152; RRID AB_390204); GFP (1:2000; Abcam, # ab13970; RRID AB_300798), TrkB (1:200; R&D AF1494; RRID AB_2155264); anti-Phox2B (1:200, kind gift of Dr. Christo Goridis, IBENS, Paris, France) and HuC/D (1:20,000, kind gift of Dr. Vanda Lennon, Mayo Clinic, MN). Sections were washed three times with PBS and incubated with secondary antibodies (Invitrogen; Jackson Immuno Research, West Grove, PA) and DAPI (Sigma, St. Louis, MO) for 1 hour at room temperature. Sections were mounted in Prolong Gold, and imaged by confocal microscopy.

### Cell quantification

#### Ganglion Cell Quantification

Ganglia were dissected from P9 to 10-month-old mice; to analyze ganglia in neonates, the upper bodies were embedded. Dissected ganglia from adults were fixed for 1.5 hours while neonate tissue was fixed for 2 hours in 4% PFA at 4° Celsius, then embedded in 100% OCT after a series of sucrose infusions. Tissue was cryosectioned at 12 μm for ganglion analysis and 14 μm for brainstem analysis. All tissue was imaged using a Lecia TCS SP8 confocal microscope and Z-stack images were obtained using Leica Application Suite Advanced Fluorescence software. All sections containing ganglia were imaged. The number of TrkB and TH positive cells in each section were counted blind as to genotype and the total number of cells composing the entire nodose-jugular ganglia (NJG), nodose-petrosal ganglia (NPG), Jugular ganglion (JG) or nodose ganglion (NG) determined. The total number of cells per ganglion, or average of left and right total number of cells if both were able to be counted, was compared between mutant and control groups using an unpaired one-tail t-test.

#### Nucleus tract solitarius (NTS) Cell Quantification

The NTS was divided into the AP region, intermediate, and caudal regions. To count Phox2b+ neurons, at least 6 equivalent planes from each region were compared using Image J. The total number of cells in each region was compared between mutants and controls using an unpaired one-tail t-test. All quantifications were done blind as to genotype.

#### Carotid body glomus cell quantification

*Wnt1-cre;Elp1* upper bodies were fixed for 2 hours at 4C and embedded in OCT as described (George et al. 2013). Consecutive sections were collected at 16 um. Slides were immunolabeled with antibody to tyrosine hydroxylase (TH). The carotid body is located in the bifurcation of the carotid artery and can be distinguished by the adjacent superior cervical ganglion. Images were taken at 63x in a single focal plan and image parameters were identical for all slides. All TH+ neurons in the carotid bodies were counted, n = 10 embryos (5 mutants and 5 controls). All quantifications were done blind as to genotype.

## Results

Mice that are null for *Elp1* are embryonic lethal because they fail to neurulate and form vasculature (Chen et al. 2009; Dietrich et al. 2011). Thus given the dual embryonic origin of visceral sensory neurons, in order to determine the requirement for Elp1 in the cranial ganglia IX and X, three different conditional knock-out mouse models were analyzed: (1) in which *Elp1* is deleted from neural crest cells but not placodes, using a *Wnt1-cre;* (Dickinson and McMahon 1992; George et al. 2013; Jackson et al. 2014); (2) in which *Elp1* is deleted from placode-derived cells and only a minority of neural crest-derived cells using a *Phox2b-cre* (Rossi et al. 2011) and (3) in which *Elp1* is deleted from nascent neurons throughout the nervous system regardless of embryonic origin using *Tuba1a-cre* (Gloster et al. 1994; Fernández et al. 2008; Ueki et al. 2016; Chaverra et al. 2017). In the new line we generated by crossing the *Phox2b-cre* driver to our floxed *Elp1* mice (George et al. 2013), *Phox2b-cre;Elp1^loxp/loxp^*, mice are born at the expected mendelian ratio of 1:4.

### Elp1 expression during visceral sensory neuron development

To determine the temporal requirement for *Elp1*, we first investigated its expression in the developing carotid body and cranial ganglia IX and X using a β-gal/LacZ reporter system (George et al. 2013) as antibodies that detect Elp1 expression with high fidelity in these tissues are not available. We found that *Elp1* is prominently expressed during development of both the nodose and petrosal ganglia (Figure 2A) and in the carotid body (Figure 2B).

**Figure 2.**
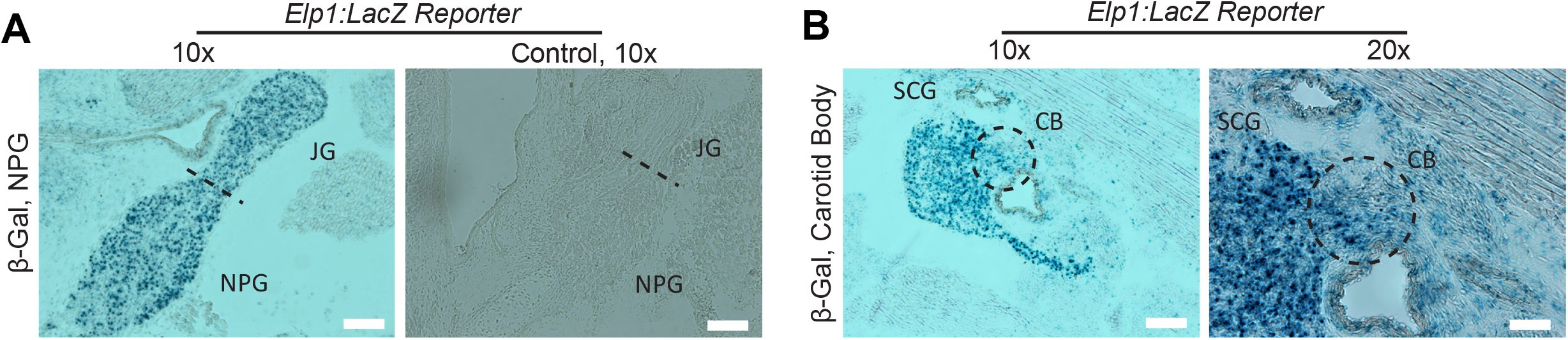
Endogenous *Elp1* expression in Nodose, petrosal and jugular ganglia and in the carotid body. *Elp1:LacZ* reporter mouse reveals β-gal staining in Elp1-expressing tissue: (A) nodose-petrosal-jugular ganglion (NPJ) complex at E17.5, with a LacZ-minus littermate shown for comparison; (B), in carotid body (CB) at E17.5, black circles surround the carotid body. Superior cervical ganglion (SCG). The cells that form the carotid body derive from cells in the SCG. (NPG): nodose petrosal ganglion, (JG): jugular ganglion. Scale bar: 50 microns (F).

To determine the activity of each Cre driver in the developing nerve IX and X ganglia, we crossed each of these lines to a GFP reporter line driven by the *Rosa26* locus, *ROSA^mT-mG^*, and imaged their expression at the end of embryogenesis, E18.5 (Figure 3). As noted by others, depending upon the age examined, it can be difficult to distinguish the boundaries between the nodose, petrosal and jugular ganglia because they are frequently fused. There is a progression in their segregation as they mature; the NPG separates into the NG and PG before P9. When all 3 ganglia appear fused, they are referred to as the “nodose-petrosal-jugular” complex (NPJ).

**Figure 3.**
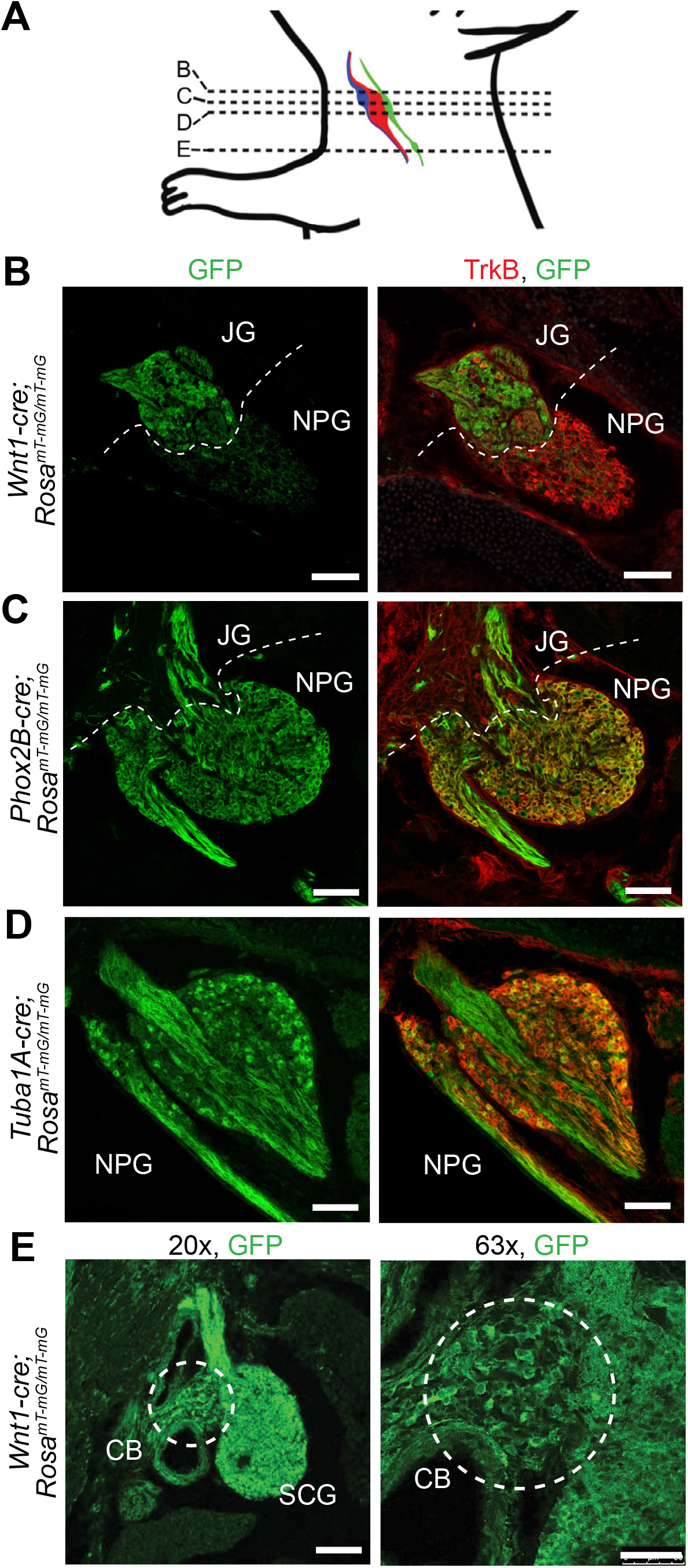
Cre activity in nodose-petrosal-jugular ganglion complex. (A) schematic of the locations of the jugular ganglion (JG, blue), nodose (red) and petrosal ganglia (green) which form the nodose petrosal complex (NPG), and carotid body (green, lower dot) is shown, with the dotted lines indicating the transverse sections locations for B, C, D, and E. *Rosa^mT-mG/mT-mG^* mice were crossed with respective Cre lines to report expression of the cre recombinase enzyme in the complex. GFP reported cre expression (left column) at E17.5 for *Wnt1-cre* (B), P1 for *Phox2b-cre* (C), and E18.5 for *Tuba1A-cre* (D). The NPG is TrkB+, while the JG is TrkB- (merged images in right column, B, C, D). Cre expression in the carotid body of E17.5 *Wnt1-cre* mice shown in (E). White dotted lines indicate ganglia borders. White dotted circle encloses CB in E.] Magnification 200x unless specified. N = 2-3 embryos/line. Abbreviations: NG: nodose ganglion, JG: jugular ganglion, PG: petrosal ganglion, NPG: nodose petrosal ganglia. Scale bar 75 microns (A-C), 100 microns (D, left), 50 microns (D, right).

We found that the *Tuba1a-cre* was expressed in about 40% of the neurons in the NPJ complex (Figure 3; data not shown), while the *Wnt1-cre* was expressed in essentially all neurons in the jugular ganglion but not in the nodose ganglion (Figure 3). The *Phox2b-cre* was expressed in essentially all neurons in the nodose-petrosal (NP) complex, (Figure 3) but only in a few cells in the jugular ganglion. These findings support previous lineage analyses of cranial ganglia and single cell RNA sequencing studies (Karpinski et al. 2016; Kupari et al. 2019). One difference we did note from the description in Rossi et al. (2011) is that a few neurons in the superior cervical ganglion (SCG) and even more in the carotid body were also GFP+ and hence expressed the Phox2b-cre (Supplementary Figure 1). In a previously published analysis of the *Wnt1-cre;Elp1* conditional knockout (cKO) we showed that the Cre was active throughout neural crest-derived gangliogenesis (George et al. 2013), but to confirm its activity in the carotid body specifically, here we asked whether the *Wnt1-cre* was expressed in the carotid body. We found robust Cre expression in the developing carotid body which was to be expected as it is neural crest derived (Figure 3).

### Placode-derived nodose/petrosal ganglion neurons require Elp1 for normal development

We and others have established that neural crest-derived neurons depend on Elp1 for survival, however whether placode-derived neurons are also dependent on Elp1 has not been determined. Since the majority of the *Wnt1-cre;Elp1* cKO mice die within 24 hrs of birth (George et al. 2013; Jackson et al. 2014), we quantified visceral sensory number just prior to birth at E18.5. Here we report that the *Phox2b-cre;Elp1* cKO new born mice also die within a few days of birth, with the longest lived mouse surviving to P18. We also found that the majority of the *Phox2b-cre;Elp1* cKO mice were smaller than their non-mutant littermates, weighed approximately 28% less than controls: at P9, the average weight of the control mice was 4.79 + 0.33 g, SEM, while the average weight of mutants was 3.44 + 0.26 g (n = 7 mutants and 8 controls; *p* = 0.007). Jackson et al (2014) also made a *Wnt1-cre;Elp1* cKO mouse which also died post-natally, which they state was due to impaired breathing associated with cleft palate. We checked whether our *Wnt1-cre;Elp1* cKO and *Phox2b-cre;Elp1* cKO mice also develop cleft palate and found that while the *Wnt1-cre;Elp1* cKO mice did show failure in palate fusion, the *Phox2b-cre;Elp1* cKO mice did not (data not shown). This finding is perhaps not surprising since *Wnt1-cre* is expressed in the cranial crest while *Phox2b-cre* is not.

Baroreceptive neurons have previously been shown to depend on the neurotrophic factors BDNF and NT-4 for their survival via TrkB activation (Erickson et al. 1996; Hellard et al. 2004). In the absence of TrkB signaling, there is a 94% reduction in nodose-petrosal cell number at birth (Erickson et al. 1996). Single cell RNAseq analysis and reporter-lineage tracking identifies TrkB as a cardinal gene expressed by the vast majority of nodose neurons (and in contrast, is only sparsely expressed in the jugular ganglion (Kupari et al. 2019; Kim et al. 2020)). To quantify changes in neuronal number in the ganglia, we used TrkB as a neuronal marker in mice just prior to or after birth (Fig. 4). We found a reduction in the number of T rkB+ neurons in the cKOs in all three mouse lines. Since the jugular ganglion is neural crest-derived and does not contain many TrkB neurons, we also quantified total neuronal number using the HuC/D antibody (kind gift of Dr. Vanda Lennon, Mayo Clinic, MN) and found a significant reduction in jugular neurons in the absence of Elp1 (Fig.4B-C).

**Figure 4.**
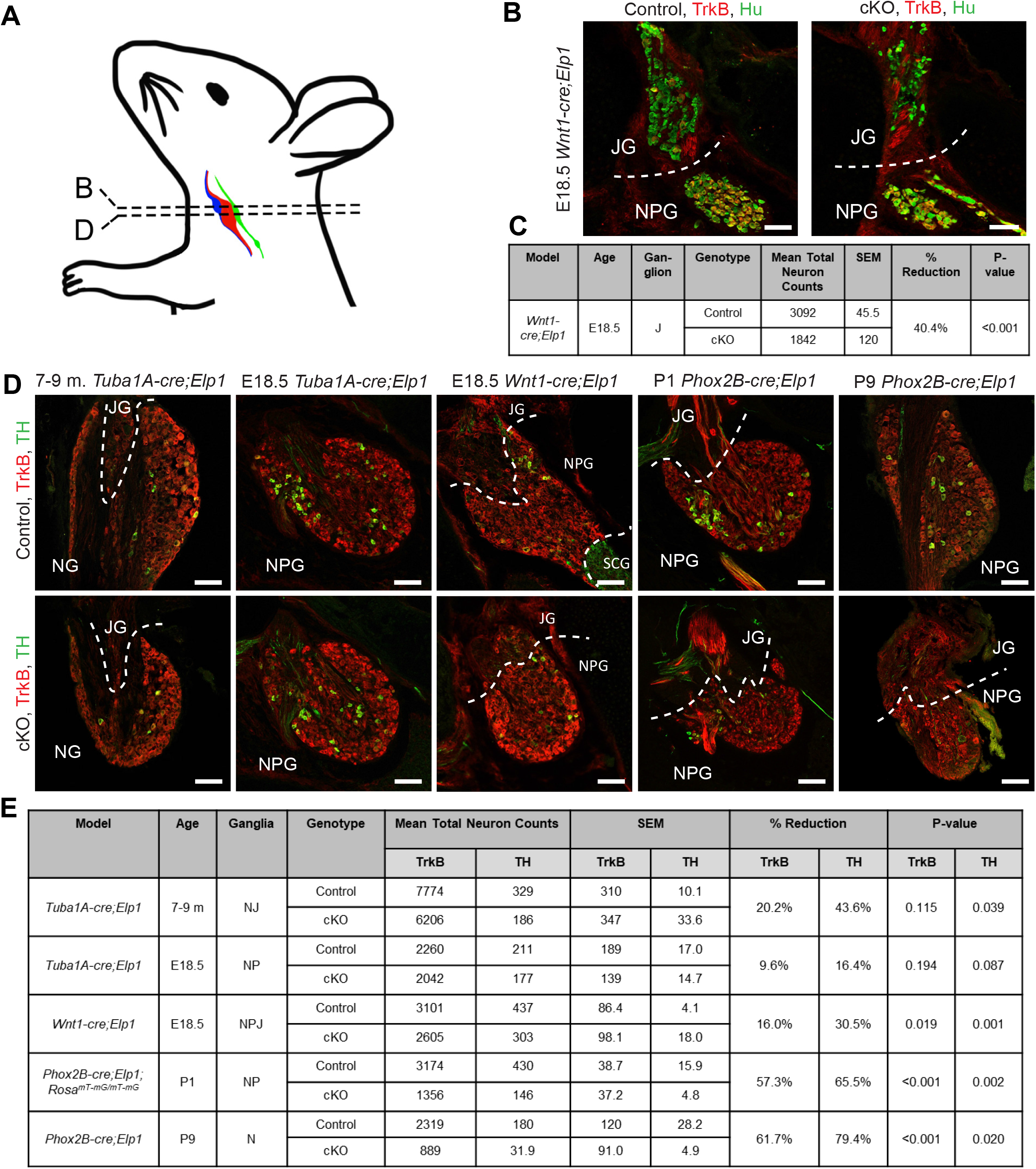
Visceral sensory neurons are reduced in the absence of Elp1. (A) Schematic depicting relative positions of nodose, petrosal and jugular ganglia. (B, C) Jugular neurons (TrkB-, HuC/D+ (green) were significantly reduced by 40% in the Wnt1-cre;Elp1 cKO mouse line. (D,E) Total ganglia cell counts (TrkB+, red) and dopaminergic neurons (TH+, green) are reduced in the absence of *Elp1* in mouse models of FD, with the NG and NPG of the *Phox2B-cre;Elp1* cKO model showing the greatest neuronal reduction compared to controls. In (E), GFP+ cells in P1 *Phox2b-cre;Elp1;Rosa^mT-mG/mT-mG^* were counted instead of TrkB+. N=4 mice for each treatment group. P-values correspond to an unpaired student’s t-test. Magnification 200x. White dotted lines denote ganglia borders. Abbreviations: JG: jugular ganglion; NG: nodose ganglion; NPG: nodose-petrosal ganglion. Scale bar = 75 microns.

Of all the mouse lines, the most dramatic neuronal loss of approximately 60% occurred in the *Phox2b-cre;Elp1* cKO (Fig.4 D-E). In contrast, in the *Wnt1-cre;Elp1* cKO, there was a reduction of only 16% of TrkB+ neurons which is consistent with the minor contribution of the neural crest to the nodose ganglion (Nassenstein et al. 2010; Karpinski et al. 2016) and/or could be due to the small number of T rkB+ (neural crest-derived) jugular neurons. We also quantified neuronal number in *Phox2b-cre;Elp1* cKO mice that were crossed to the *ROSA^mT-mG^* reporter and found again about a 60% reduction in neurons. In this case, the number of (Cre+) GFP+ cells were counted since as we showed previously, the vast majority of them (>90%) are TrkB+. This reduction is consistent with the visible decrease in diameter of the vagus nerve at P9 in the *Phox2b-cre;Elp1* cKO compared to control (Supp Figure 2). There were no significant differences between left and right ganglia nor differences between males and females. The number of NPJ neurons in the *Tuba1a-cre* at E18.5 was not significantly reduced in the mutants. Since of these lines, only the *Tuba1a-cre;Elp1* cKO mice live into adulthood, we also quantified TrkB+ neurons in their adult NJ complex and found approximately a 20% reduction. While the difference was insignificant, it was increased in the adult compared to the E18.5 ganglia suggestive of a progressive decline. These data are perhaps not surprising given that the *Tuba1a-cre* is only expressed in about 40% of NPJ neurons..

### Dopaminergic neurons in the Nodose ganglion depend on Elp1 for development

We also investigated the requirement for *Elp1* in the dopaminergic neurons located in the NP complex because they have been shown to innervate the chremosensory glomus cells in the carotid body (Katz et al. 1983; Katz and Black 1986) and are required for normal respiratory drive (Erickson et al. 1996). These cells are known to give rise to the super laryngeal nerve (Kummer et al., 1993) which detects sensory stimuli for the swallowing reflex which is also severely impaired in FD patients (Palma et al., 2014) and innervates the esophagus and stomach (Kummer et al. 1993). As shown in Figure 4, the loss of dopaminergic neurons – as marked by expression of tyrosine hydroxylase (TH), was more dramatic than that of the reduction in TrkB+ neurons, being reduced from 16-80% depending on the mouse line and age. The biggest reductions occurred in the *Phox2B-cre;Elp1* cKO mouse, suggesting that the majority of the TH+ neurons are placode derived. In contrast, only about 2% of TH+ neurons were located in the (neural crest-derived) jugular ganglion in all of the mouse models analyzed. In the *Tuba1a-cre;Elp1* cKO, the reduction in TH+ neurons was significant in the older adult mice (44%), but not at E18.5 (16%), consistent with a progressive loss of these neurons as the mice age.

Interestingly, the TH+ neurons were typically found in clusters within the NP complex (Fig. 4A and 5A), which could suggest a clonal origin of these neurons from a common precursor cell(s). TH clusters were present in both mutants and controls in all three mouse lines. Co-localization analysis of *Phox2b-cre;Elp1;Rosa^mTmG^* tissue sections stained with TH antibody, show that the TH+ neurons in the Nodose-petrosal ganglion do express Phox2b and are targeted by the Phox2b-cre (Fig. 5). Quantification of the number of TH+ cells per cluster revealed a significant reduction in the cKO mice compared to control littermates in all three lines (Fig. 5). There is conflicting information on whether TH clusters are localized to the nodose or petrosal ganglia specifically (Kummer et al. 1993) but since they were found in post-natal and adult mice at times in which the nodose and petrosal ganglia can be clearly distinguished from each other and selectively dissected, we conclude they are clearly present in the nodose ganglion.

**Figure 5.**
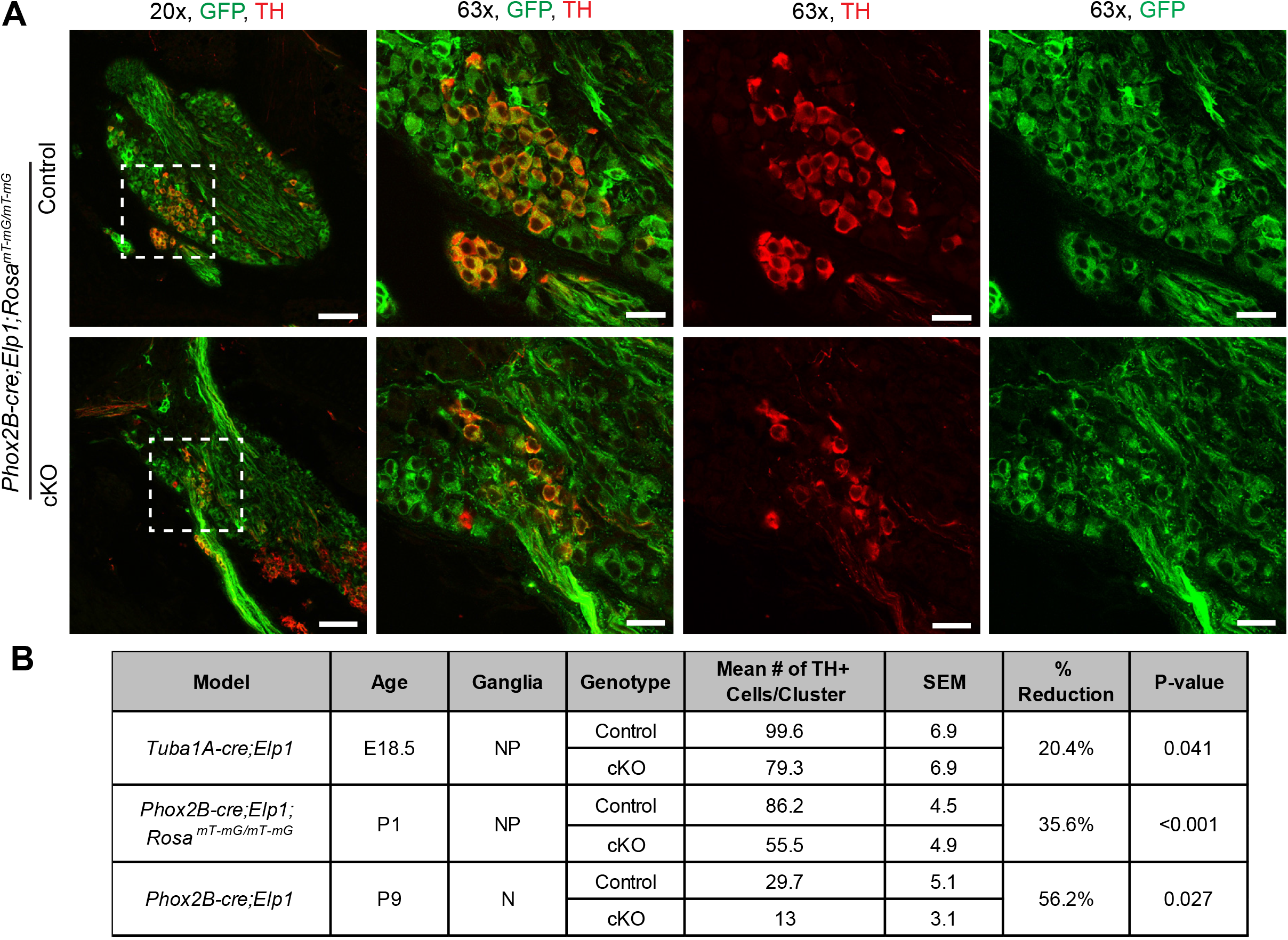
TH+ cell clusters in the NPG express Phox2b-cre and are reduced in number with deletion of *Elp1*. TH+ clusters in the nodose petrosal ganglia (NP) are Phox2B+ as shown by colocalization of GFP and TH in P1 *Phox2b-cre;Elp1;Rosa^mT-mG/mT-mG^* mice (A, 20x and 63x). White dashed box depicts the location of the 63x image. The number of TH+ cells in the NPG cluster are significantly reduced in cKO *Phox2b-cre;Elp1^LoxP/LoxP^;Rosa^mT-mG/mT-mG^* mice compared to control *Phox2b-cre;Elp1^+/LoxP^;Rosa^mT-mG/mT-mG^* mice at P1 (A, B). Number of TH+ cells in the cluster are also reduced in the nodose ganglion (N) of P9 *Phox2b-cre;Elp1* cKO compared to control and the NP of E18.5 *Tuba1A-cre;Elp1* cKO compared to control (B). N=4 for each group of mice. Scale bar at 20x = 75 microns; at 63x = 25 microns.

### Chemoreceptive glomus cells in the carotid body require Elp1 for normal development

To determine whether neural crest-derived glomus cells also require *Elp1* for their development, the number of glomus cells - as identified morphologically, by their stereotyped location between the external and internal carotid, and by their expression of tyrosine hydroxylase - was quantified in the *Wnt1-cre;Elp1* cKO and control littermates at the end of embryogenesis (E18.5). The cKO carotid body was dramatically reduced in size compared to control littermates and the number of TH+ glomus cells was significantly reduced by approximately 40% in the cKO carotid bodies (Figure 6 A, B) compared to controls.

**Figure 6.**
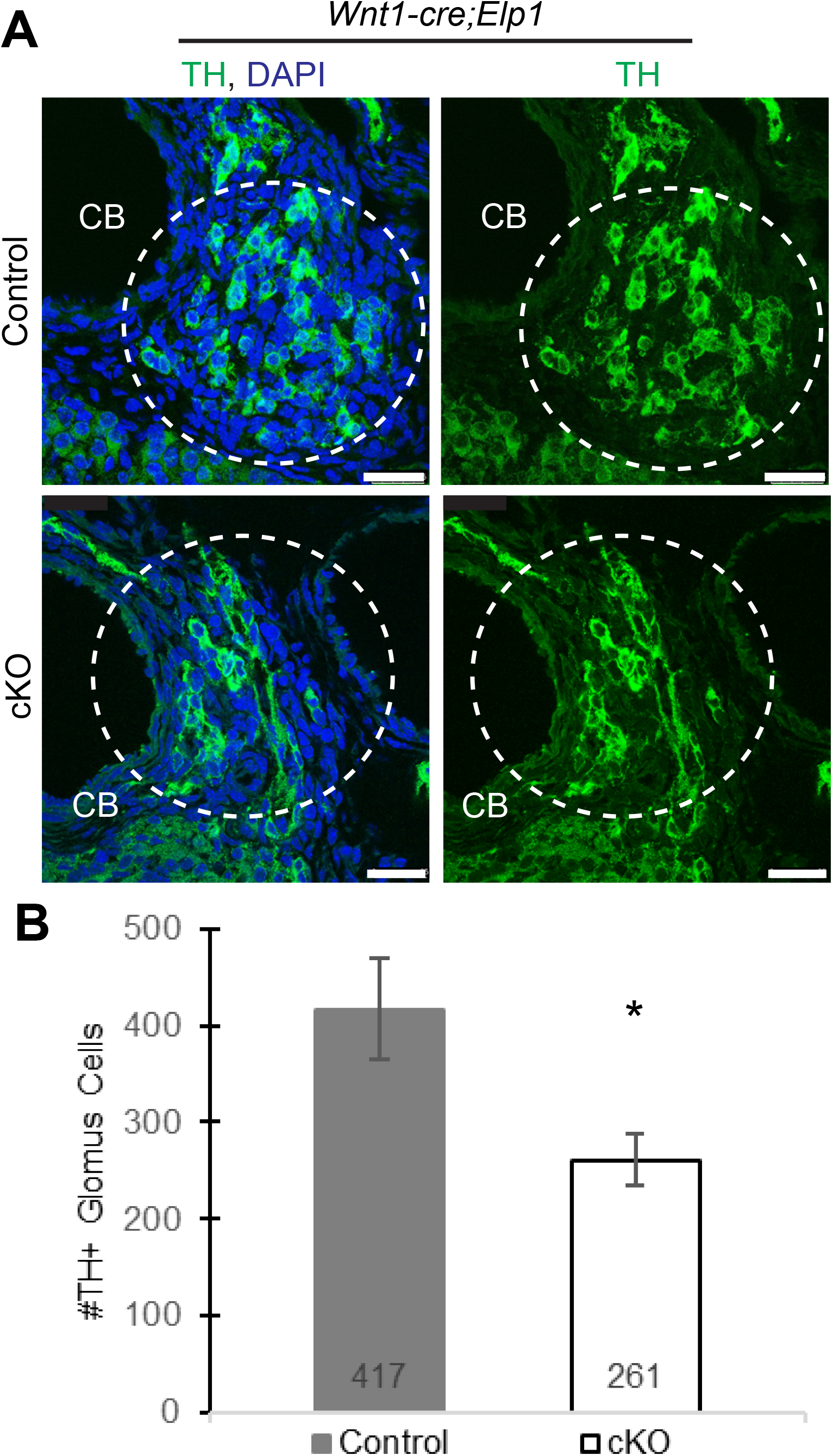
Glomus cells in the carotid body require *Elp1* for normal development. (A-B) Embryonic E18.5 *Wnt1-cre;Elp1* transverse sections were stained with antibodies to tyrosine hydroxylase (TH), and the number of glomus cells was quantified through each carotid body. N=5 mice per condition. P-value, <0.026, unpaired student’s t-test. (CB), Carotid body. Scale bar: 25 microns

### Brainstem nuclei that process visceral sensory information depend on Elp1

Remarkably, the peripheral visceral sensory neurons and their central targets in the brainstem not only all express Phox2b (Tiveron et al. 1996; Dauger et al. 2003) but Phox2b specifies the entire neural circuitry that mediates both the baroreflex and chemoreflex (Dauger et al. 2003). Deletion of *Phox2b* impairs development of the sensory afferents in the carotid body, petrosal and nodose ganglia, their brainstem targets in the NTS and efferent arms for parasympathetic and enteric circuits. Consequently, we used the *Phox2b-cre;Elp1^loxp/loxp^* mouse line to investigate whether the central components of respiratory and cardiovascular circuits were also dependent on *Elp1* expression.

Brainstems were sectioned from the pons to the medulla and immunolabeled with antibodies to Phox2b (Figure 7). Reduced numbers of Phox2b+ neurons were evident throughout the brainstem of the *Elp1* cKO mouse compared to control littermates. The number of Phox2b+ neurons in the central targets of the NPJ axons were quantified in the area postrema (AP), nucleus tract solitarius (NTS), ventral respiratory group (VRG) and nucleus ambiguous (NA) in the *Phox2b-cre;Elp1* cKO and littermate controls at P0. The intermediate region of the NTS receives input from vagal gastrointestinal afferents, while the caudal NTS receives input from the vagal and glossopharyngeal baroreceptors, vagal chemoreceptors, vagal cardiac receptors, and vagal pulmonary receptors (Cutsforth-Gregory and Benarroch 2017). The VRG (ventral respiratory group) and RTN (retrotrapezoid nucleus) are functionally interconnected in response to chemoreceptor drive and the RTN neurons are themselves central CO2 chemoreceptors (Ott et al. 2012)Souza et al., 2019). Interestingly, there are numerous Phox2b+/GFP-neurons in adjoining nuclei that are missing in the cKO brainstem, suggesting they are lost indirectly. As shown in Figure 7, the numbers of Phox2b+ neurons in all of these nuclei were reduced by approximately 50% in the cKO compared to their littermate controls.

**Figure 7.**
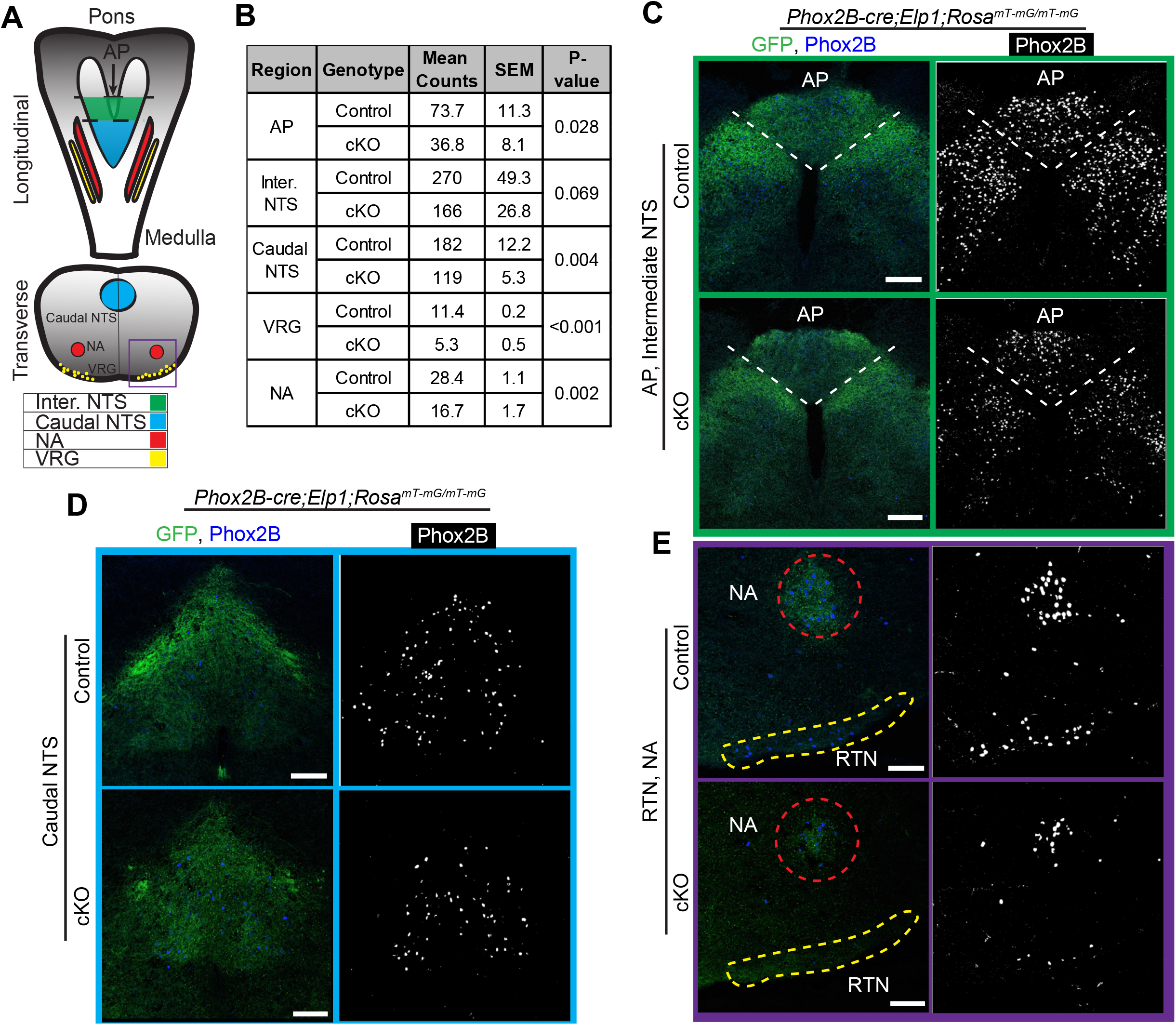
Neurons in the Brainstem respiratory circuitry are reduced in the absence of *Elp1.* Brainstem regions analyzed are schematized in (A). Phox2b+ neurons in the brainstem respiratory nuclei were visualized with a Phox2b antibody, and shown to be significantly reduced in the AP, NTS, VRG, and NA of cKO *Phox2B;Elp1^LoxP/LoxP^;Rosa^mT-mG/mT-mG^* mice compared to control *Phox2b;Elp1^+/LoxP^;Rosa^mT-mG/mT-mG^* mice at E18.5 or P1 (B). The intensity of GFP staining, marking Phox2b-cre expressing cells, is also reduced in each region (C, D, E). N=3 mice for each analysis. Abbreviations: AP: area postrema, NTS: nucleus of the solitary tract, VRG: ventral respiratory group, NA: nucleus ambiguous, RTN: retrotrapezoid nucleus. Scale bar = 75 microns.

### Elp1 is required for normal innervation of baroreceptive peripheral targets

For sensory neurons to function effectively, they must innervate their peripheral targets successfully. Previous studies have established that neural crest-derived sensory neurons that lack *Elp1* fail to normally innervate their peripheral targets (George et al. 2013; Jackson et al. 2014; Ohlen et al. 2017); however, whether placode-derived sensory neurons also require *Elp1* for peripheral target innervation has not been investigated. Baroreceptive neurons in the nodose ganglion innervate the carotid sinus and aortic arch (Min et al. 2019) by extending decorated “claws” that circumscribe the diameter of the aortic arch. Via activation of mechanoreceptive proteins such as Piezo 1 and 2, baroreceptive neurons are stimulated by changes in blood pressure in these vessels (Zeng et al. 2018; Min et al. 2019). Thus in their absence, changes in blood pressure can’t be detected nor relayed to the CNS to appropriately adjust cardiac output. To determine if the baroreceptive targets in the aortic arch and subclavian artery were innervated normally in the *Phox2b-cre;Elp1* cKO mice, the *Rosa^mTmG^* reporter line was crossed into this line, aortic arches and subclavian arteries dissected, and whole mount staining with GFP antibodies conducted (Figure 8, Supplemental Figure 3). Z-stacks were collected through the entire aortic arch, from the anterior to the posterior sides of the arch, thus revealing both the ligament and saddle regions of the aortic arch. The left nodose innervates the aortic arch and right nodose innervates the right subclavian artery. Vagal afferents also innervate the right subclavian artery at the bifurcation of the brachiocephalic artery. The aortic depressor nerve (ADN) branches from the vagus. Near the aortic arch, the left ADN bifurcates with one branch forming the saddle region on the posterior side of the arch and the other branch forming the ligament region on the anterior side on the arch (A, Tomato shows aortic arch). The right baroreceptors form a claw around the right subclavian artery (RSA; B, tomato). Innervation of the aortic arch (Figure 8B) in three mutants and controls, and in the RSA in four mutant and controls, (Figure 8C) all reveal severe reductions in innervation in the cKO compared to littermate controls, with some variation amongst the mice. The data show that the right and the left baroreceptor branching in both the saddle and the ligament are decreased in density in cKO mice compared to their control littermates, with the left ADN appearing thinner than that in controls. Afferents terminated in two types of endings, the flower-sprays and end-nets as described (Cheng et al. 1997; Min et al. 2019). Both types of endings are reduced in the cKO mice (Supplemental Figure 3), with the end-net endings severely reduced in the ligament. Interestingly, many of the axons terminating in end net endings appear to be misrouted and extend to regions ectopic to their normal target in the ligament (Fig. 8B, white arrowheads). Thus although the viscerosensory axons that comprise the ADN navigate successfully to and invade the aortic arch and RSA, the final innervation of target tissue is impaired for the majority of the baroreceptive axons in the absence of Elp1.

**Figure 8.**
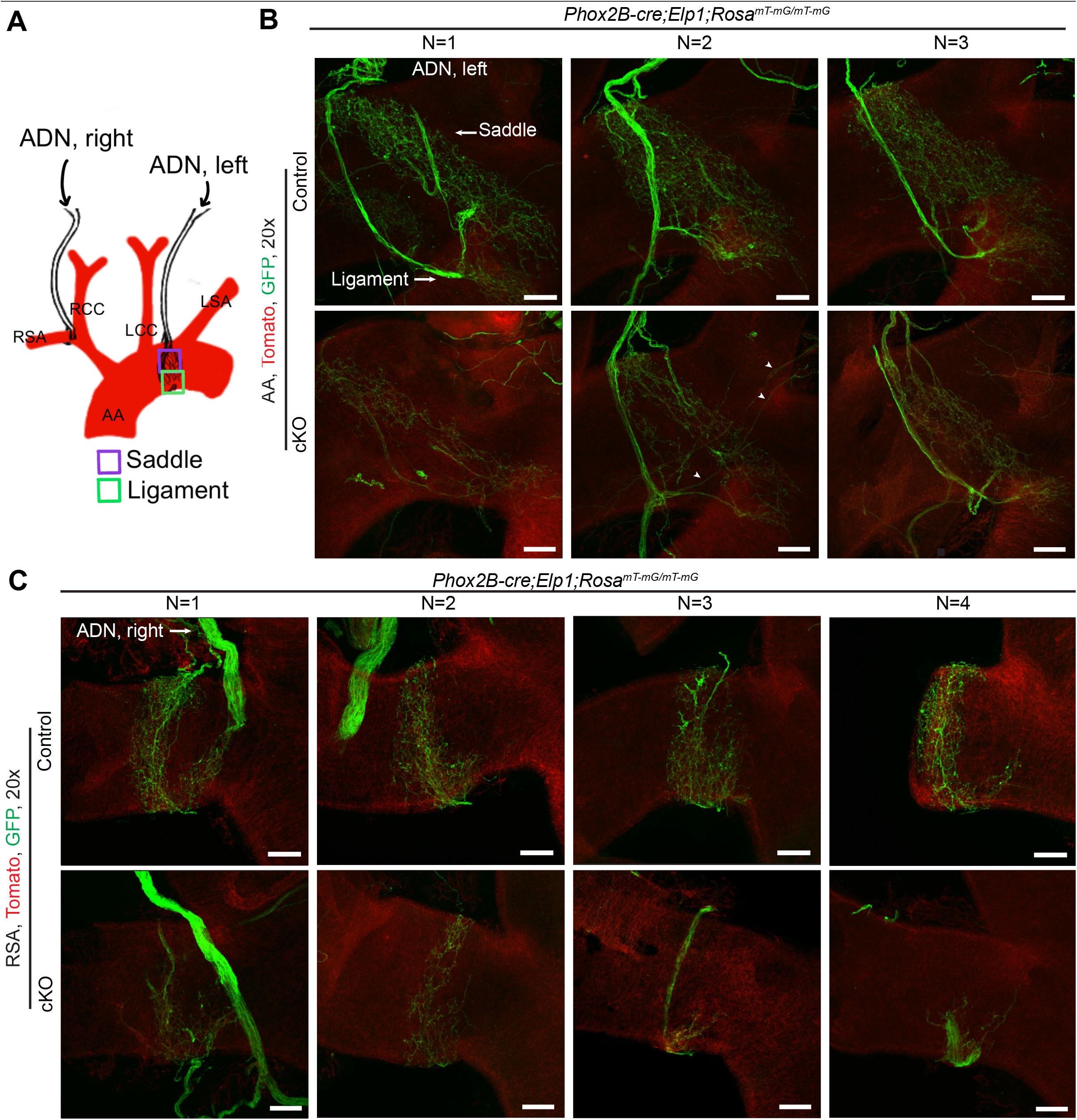
Elp1 is required for normal innervation of the aortic arch. (A) Schematic depicting visceral sensory neurons innervation of the saddle and ligament regions of the aortic arch (B) and the right subclavian artery (C), at the bifurcation of the brachiocephalic artery. Right and baroreceptive endings in both the saddle and the ligament are decreased in density and branching in cKO *Phox2b;Elp1^LoxP/LoxP^;Rosa^mT-mG/mT-mG^* mice compared to control *Phox2b;Elp1^+/LoxP^;Rosa^mT-mG/mT-mG^* mice. N = 3 - 4 mice per condition. White arrowheads in (B) point to ligament axons that rather than staying within the ligament, grow in an incorrect direction to traverse through regions of the aortic arch they don’t normally grow into. AA: aortic arch, RSA: right subclavian artery, ADN: aortic depressor nerve. Scale bar = 100 microns (B), 75 microns (C).

## Discussion

Failure in key cardiovascular and respiratory reflexes such as the baroreflex and chemoreflexes are clinical hallmarks of FD, with often fatal consequences. Using mouse models for FD, this study delineates the potential underlying cellular basis for these failed reflexes. While impaired somatic sensory neuron development has been demonstrated in several FD mouse model studies, here we report the first investigation of visceral sensory neurons, which mediate these critical cardio-respiratory reflexes. Our data indicate that despite arising from a distinct embryological lineage, visceral sensory neurons, like somatic sensory neurons, also depend on *Elp1* for development. While the neural crest-derived somatic sensory neurons in the dorsal root ganglia (DRG) mediate somatic functions such as pain, temperature, pressure and proprioception, the cranial-placode derived visceral sensory neurons located in the vagal and glossopharyngeal ganglia are vital for key autonomic functions such as respiration, cardiovascular regulation, and gastrointestinal function. Moreover, this study reveals that *Elp1* is required in the entire CNS/PNS circuitry underlying baroreception and chemoreception: in both the central brainstem targets of the visceral afferents, and, for proper innervation of their baroreceptive peripheral targets.

The fundamental importance of Elp1 in development is apparent by the fact that mice that are null for *Elp1* die by E10, due to failures in neurulation and vasculature formation (Chen et al. 2009; Dietrich et al. 2011). Yet why dependence on *Elp1* varies temporally and by cell type is not understood. This study expands our knowledge of the cell types that require *Elp1* for their development since in spite of being broadly expressed in both the CNS and PNS, not all neuronal populations depend on *Elp1* for their development and survival (George et al. 2013; Ueki et al. 2016). For example, in the DRG, TrkA and TrkB+ neurons require *Elp1* during development, but TrkC+ neurons do not. Yet, in adulthood, the TrkC subpopulation which constitute proprioceptors, do progressively die causing the degenerative gait ataxia of FD patients and mice (Morini et al. 2016). Similarly, our work has shown that in spite of *Elp1’s* robust expression in the retina, retinal neurons do not require *Elp1* for their development, however, within 2 weeks of birth, *Elp1* cKO retinal ganglion cells progressively die, while strikingly, the other retinal cell types can persist and function in the absence of any Elp1 expression. Here we add visceral sensory neurons to the list of neurons that do require Elp1 for their development and survival.

Elp1 functions with the five other elongator subunits, Elp2-Elp6, to form the Elongator complex which modifies tRNA to facilitate the translation of mRNAs that exhibit a biased usage of either AA- or AG-ending codons for lysine, glutamine, and glutamic acid (Huang et al. 2005; Bauer and Hermand 2012; Goffena et al. 2018). It is possible then that the codon-usage of a particular cell type’s transcriptome could determine to what extent an Elp1-deficient cell lives or dies. Moreover, the fact that mutations and/or variants in *ELP2-4*, are associated with distinct neurological disorders (albeit with some overlapping features) ranging from intellectual disability to ALS but excluding the sympathetic, sensory, and optic neuropathies exhibited by FD patients, may speak to variation in function of the Elp subunits independent of their joint actions as part of the elongator complex. Although the vast majority of Elp1 is localized to the cytoplasm, there are reports that the elongator complex is also involved in transcriptional elongation (Otero et al. 1999; Hawkes et al. 2002; Close et al. 2006; Morini et al. 2021); this effect may be the indirect consequence of changes in the cells’ proteome that occur with loss of elongator function.

The visceral sensory neurons in the nodose ganglion are generated and specified in cranial placodes and upon becoming post-mitotic, they delaminate as specified, differentiating neurons and migrate to their targets (Graham et al. 2007; Blentic et al. 2011; Moody and LaMantia 2015). Neurogenesis does not occur in the ganglion (Blentic et al. 2011). This is in contrast to how neural crest cells give rise to neurons: neural crest cells migrate to multiple locations in the head and trunk and only upon stopping to colonize their targets, do they either differentiate into neurons directly or give rise to progenitor cells that then differentiate into neurons (Maro et al. 2004; George et al. 2007; George et al. 2010; Ventéo et al. 2019). For example, in the DRG, TrkA+ neurons derive from intermediate Pax3+ progenitors (George et al. 2010; George et al. 2013) that are especially sensitive to reductions in Elp1. While here we report that Elp1 is expressed in both the NP ganglia and in the carotid body during development, we did not determine at what developmental stage the reduction in the glomus cells and NP neurons in the cKO occurred. If in fact, Elp1 is required in nodose neurons while still in their placode, this study would have missed that developmental window since Phox2b is first expressed when neurons are differentiating in the nodose ganglion.

What our data do reveal is that *Elp1* is required to acquire a full complement of TrkB+ neurons by birth. NP neurons, in particular those that mediate the afferent pathway driving respiration and baroreception, have been demonstrated to depend on TrkB activation via BDNF and NT4 for their survival (Erickson et al. 1996). Mice in which *BDNF* was deleted have dysfunctional respiratory behavior including a reduction in chemosensory drive. At a cellular level, 94% of NP neurons are missing in TrkB knock-outs, vs. approximately a 50% loss if either BDNF or NT-4 is deleted (Erickson et al. 1996). Here in the *Phox2b-Elp1* cKO we lost about 60% of the TrkB+ neurons, potentially consistent with an overlap in the TrkB signaling pathway. Interestingly, patients with Rett syndrome have impaired autonomic respiratory function (Katz et al. 2009) which in mouse models is marked by reductions in nodose neurons and can be ameliorated with TrkB agonists (Schmid et al. 2012; Kron et al. 2014). Given the dependency of nodose ganglion neurons on BDNF signaling, and studies showing that Elp1 cKO neurons have impaired NGF retrograde transport and TrkA signaling (Abashidze et al. 2014; Naftelberg et al. 2016; Li et al. 2020) it will be interesting to investigate in future studies whether nodose neurons in FD die due to faulty TrkB retrograde signaling.

To investigate the dependency of the visceral sensory neurons on Elp1, we used three different conditional knock-out lines. As described previously, since the NPJ complex includes cells that derive from both the neural crest and from cranial placodes, to ensure each cell population was probed, we used a *Wnt1-cre* to target the neural crest cell derivatives, the *Phox2b-cre* to target the nodose and inferior petrosal neurons, and a *Tuba1a-cre* which targets about 40% of neurons throughout the NPJ complex. In all lines, the *cre* is expressed during development (Figure 3), and in all three lines, neuronal number was reduced in the nodose, NP or NPJ complex. The largest deficit was in the *Phox2b-cre;Elp1^loxp/loxp^* line which could be due to the high fidelity and robust expression of the *Cre* (Figure 3) in the nodose ganglion. Interestingly though, in spite of its ubiquitous expression in TrkB+ neurons, deletion of *Elp1* does not result in the loss of all TrkB+ or of all TH+ neurons by birth. Moreover, here we show that loss of Elp1 reduced glomus cell number and jugular neuronal number by 40%. Similar results have been discovered in other studies: deletion of *Elp1* in the neural crest causes the loss of 40% of DRG and of about 70% of sympathetic neurons (George et al. 2013; Jackson et al. 2014). This is a clinically promising finding since if upheld in humans, therapeutic interventions that protect the remaining neurons could be feasible in FD infants.

Within the NP complex, one of the most functionally important cell populations are the dopaminergic neurons which innervate the carotid body and transmit key sensory information on levels of oxygen, carbon dioxide and pH, i.e. chemoreceptive information, to the brainstem (Erickson et al. 2001). We found that this subpopulation is particularly dependent on Elp1 expression and were reduced in number from 30 – 65% depending on the mouse line. Kummer et al 1993 found that TH+ nodose ganglion neurons project axons to the esophagus and stomach, while TH+ petrosal ganglion neurons project to the carotid body (Katz and Black 1986; Kummer et al. 1993). Many of the TH+ neurons appear in clusters that reside in the nodose ganglion near the exit of the superior laryngeal nerve (Kummer et al. 1993) and were retrogradely labeled by injections of tracers into the esophagus and stomach. A reduction in this cell population most likely contributes to the severe swallowing impairment suffered by FD patients in addition to the impaired response to hypoxia and hypercapnea in FD patients.

Here we show that deletion of *Elp1* in the neural crest lineage resulted in approximately a 40% reduction in glomus cells by E17.5 (Figure 6) and the carotid body itself was reduced in size in these cKOs. The carotid glomus cells derive from the superior cervical ganglion (SCG; (Kameda 2005; Kameda et al. 2008)) and we and others have shown that at the ages we examined (E17.5-E18.5), neuronal number in the SCG is reduced by approximately 70% (George et al. 2013; Jackson et al. 2014) in *Elp1* cKO mice, hence it is not surprising that loss of *Elp1* would consequentially reduce the number of glomus cells in the carotid body. This finding of reduced glomus cell number may explain the blunted chemoreflex responses to hypoxic conditions in FD patients (Palma et al. 2019). Hence loss of *Elp1* could interfere with this sensory arm both autonomously in the glomus cells, and indirectly via reduced innervation of glomus cells by the dopaminergic afferents in the NP complex (Conover et al. 1995). Interestingly, recent work has expanded our understanding of the chemosensory role of this organ: in addition to detecting changes in oxygen, CO2 and pH, glomus cells are also responsive to changes in blood levels of glucose, lactate, insulin and leptin (Ortega-Sáenz and López-Barneo 2020). The glial sustentacular cells can act as stem cells and produce new glomus cells under chronic hypoxemic conditions (Pardal et al. 2007). It would be interesting to know if they are triggered to generate new glomus cells over the life span of FD patients.

Given that previous studies (Abashidze et al. 2014; Jackson et al. 2014; Ohlen et al. 2017) revealed failures in peripheral target innervation by Elp1 cKO sensory neurons, we investigated the integrity of innervation of baroreceptive targets. Here we show that while many vagal sensory axons do reach the aortic arch and right subclavian artery, they do not elaborate the normal complex array of endings neither in the saddle nor along the ligament (Figure 8). This incomplete target invasion could be the result of impaired retrograde transport of neurotrophins from their targets (Naftelberg et al. 2016). Alternatively, axonal branching has been shown to be dependent on spatial and temporal localization of mitochondria at branch points in order for tubulin to invade the branches (Smith and Gallo 2018), a process which is impaired in Elp1 cKO neurons (Ohlen et al. 2017). Misrouted ligament axons were also detected suggesting that some axonal guidance mechanisms in the aorta also depend on Elp1 signaling. Taken together, impaired arterial innervation could significantly negatively impact the normal sensory response to elevated or decreased arterial pressure.

In addition to identifying the potential cellular basis for the failure in the sensory arm of the baroreflex in FD, we also demonstrate here that the central circuitry that integrates this sensory input is marked by significant reductions in cell number. An elegant series of studies by Brunet, Goridis and their colleagues demonstrated that Phox2b remarkably delineates the entire cardiorespiratory reflex pathway (Pattyn et al. 1999; Dauger et al. 2003). By immunolabeling the brainstems of *Phox2b-cre;Elp1* cKO mice with antibodies to Phox2b, we found that the target nuclei for the NP afferents in both the intermediate and caudal regions of the NTS were significantly reduced by approximately 40% in the Elp1 cKO mice by E18.5 (Figure 7). Neurons in the NTS then transmit this sensory input information to neurons in the nucleus ambiguous which were reduced by about 50% in the Elp1 cKO. Similar reductions were found in the ventrally-located medullary relay nuclei including the RTN which conveys information regarding CO2 levels as part of the central respiratory chemoreflex (Souza et al. 2019). While examining the Phox2b+ neurons in the brainstem, it became evident that there were also fewer neurons located in the area postrema (AP); upon quantification, a significant reduction of about 50% of the AP neurons was detected. Chemoreceptor afferents that innervate the carotid body have also been shown to project to the AP in addition to their major projection to the NTS (Finley and Katz 1992). Finding reductions in specific brainstem nuclei that mediate cardiorespiratory reflexes may help explain the cellular basis for the major cause of death of FD patients, Sudden death during sleep (SUDS) (Palma et al. 2017; Palma et al. 2019). The impaired ventilatory responses of FD patients to hypoxia and hypercapnia can be particularly dangerous while patients sleep and underlies the sleep-disordered breathing experienced by 95% of patients (Palma et al. 2019). We show here both reduced numbers of the chemoreceptors in the carotid body, but also reduced cell number in the NP ganglia, in addition to reduced neuronal number in the brainstem respiratory circuitry. In a previous analysis we conducted on the brainstem in the *Tuba1a-cre;Elp1* cKO mice, we not only found widespread expression of Elp1 throughout the brainstem, but also discovered a reduced number of cholinergic neurons in the dorsal motor nucleus of the vagus (Chaverra et al. 2017). These findings also are supported by autopsy studies which found brainstem atrophy in FD patients at death and dysfunctional brainstem reflexes in patients (Cohen and Solomon 1955; Brown et al. 1964; Gutiérrez et al. 2015). Collectively our data may explain the cellular basis for the patient chemoreflex failure which underlies significant patient morbidities.

### Summary

In summary, the data reported here indicate that *Elp1* is essential both in placode and neural crest derived neurons and that it exerts comparable effects including survival, axonal morphology and target innervation in both lineages. Given the important surveillance role that vagal sensory neurons exert in maintaining homeostasis, and their diversity in terms of function and gene expression, future experiments should include single cell RNA sequencing studies to identify the resilient neurons vs those that are lost in the *Elp1* cKO vagal ganglia. Such an approach would not only help us understand differential cell dependence on *Elp1*, but also inform strategies for therapeutic interventions.

---

**Supplementary Figure 1.** Low magnification view of location of visceral cranial ganglia IX and X, carotid body and Superior cervical ganglion and expression of the *Phox2b-cre.* A schematic of the locations of the jugular ganglion (JG, blue), nodose ganglion (NG, red), petrosal ganglion (PG, green), carotid body (CB, purple), and superior cervical ganglion (SCG, black) is shown (A). Mice that are heterozygous for *Elp1^loxp^* (i.e. express no mutant phenotype because they have one normal copy of *Elp1)* and express *Phox2b-cre* and *Rosa^mT-mG/mT-mG^* were sectioned at P1 and stained with antibodies to TrkB and TH. The sagittal sections depicted in (B) show that the JG is GFP- and TrkB-, the NG is GFP+ and TrkB+, while the CB and SCG are TrkB- but do contain some GFP+ cells, meaning that some cells in the CB and in the SCG do express the *Phox2b-cre*. GFP+ NP axons can be detected running through the JG, but the neurons in the JG itself do not express GFP (lower panels). White dotted lines indicate separations of ganglia or separation of CB from SCG. (C) Colocalization of TH and GFP in the CB and SCG are shown with white arrows pointing to cells that co-express TH and GFP. N = 2 mice. Abbreviations: CB, carotid body; PG, petrosal ganglion; NP, nodose-petrosal ganglia; SCG, superior cervical ganglion; NG, nodose ganglion; JG, jugular ganglion. Scale bar = 250 microns (B); 25 microns (C).

**Supplementary Figure 2**: Vagus nerve is reduced in diameter in *Elp1* cKO mice compared to control littermates. (A) A schematic identifying the location for analysis of vagus and hypoglossial nerves in (B). *Phox2b-cre;Elp1* control and cKO mice at P9 were dissected, and carotid arteries removed to reveal their vagus nerves (black arrows). Their hypoglossal nerves (indicated by the white arrow) are comparable in diameter between the cKO and control, but the vagus nerve in the cKO mouse is smaller in diameter than it is in the control mouse.

**Supplementary Figure 3. (**A) Schematic depicting the locations of the left baroreceptors, with the saddle (purple) and the ligament (green) indicated. End-net endings (red arrows) and flower spray endings (yellow arrows) for the baroreceptors in the saddle (purple border, upper panels) and the ligament (green border, lower panels) of cKO *Phox2b;Elp1^LoxP/LoxP^;Rosa^mT-mG/mT-mG^* mice are reduced compared to control *Phox2b;Elp1^+/LoxP^;Rosa^mT-mG/mT-mG^* mice (B). N=4 mice from each condition. Abbreviations: AA: Aortic Arch, LCC: left common carotid, LSA: left subclavian artery. Scale bar = 25 microns.

## Acknowledgements

This work was funded by NIH R01DK117473 and R01NS086796.

