## Supplementary figures and images for "Elp1 is required for development of visceral sensory peripheral and central circuitry"

### Supplemental data

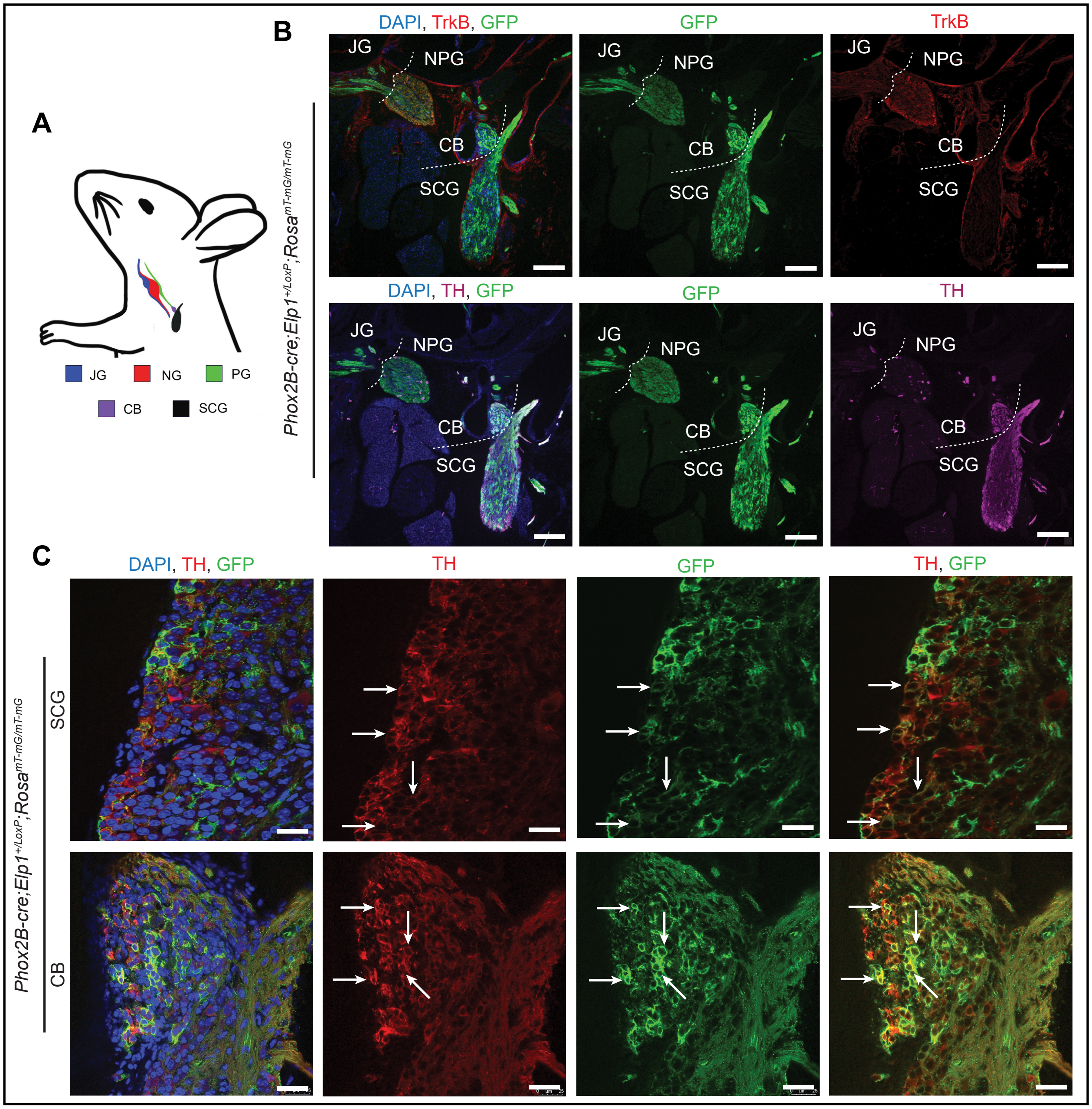


Supplementary Figure 1

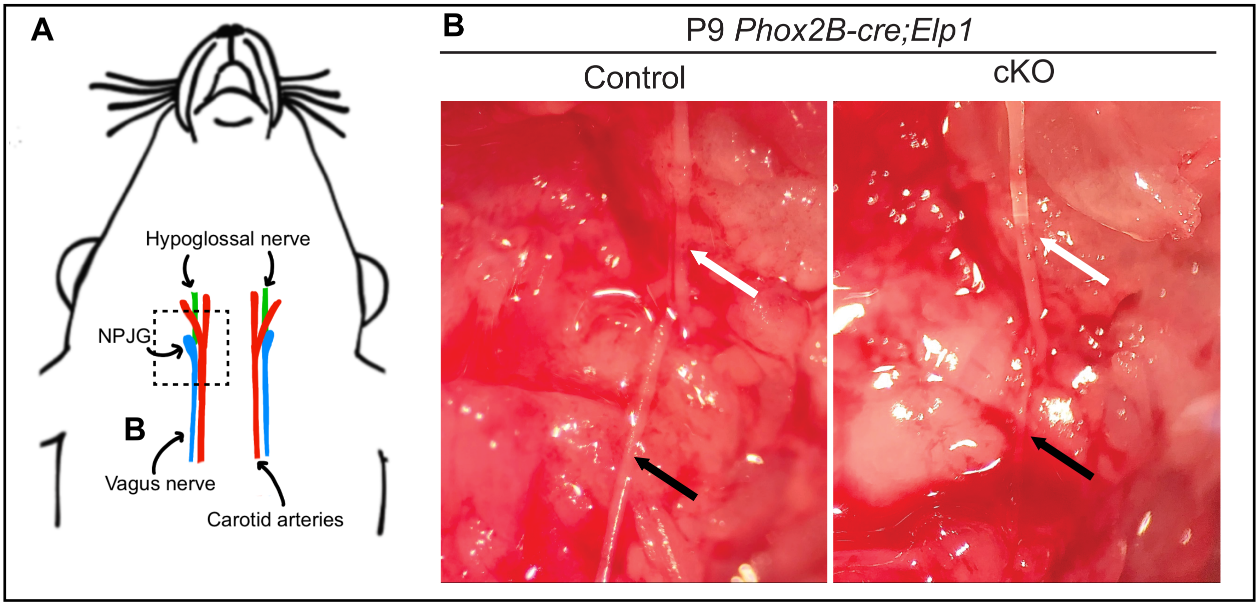


Supplementary Figure 2

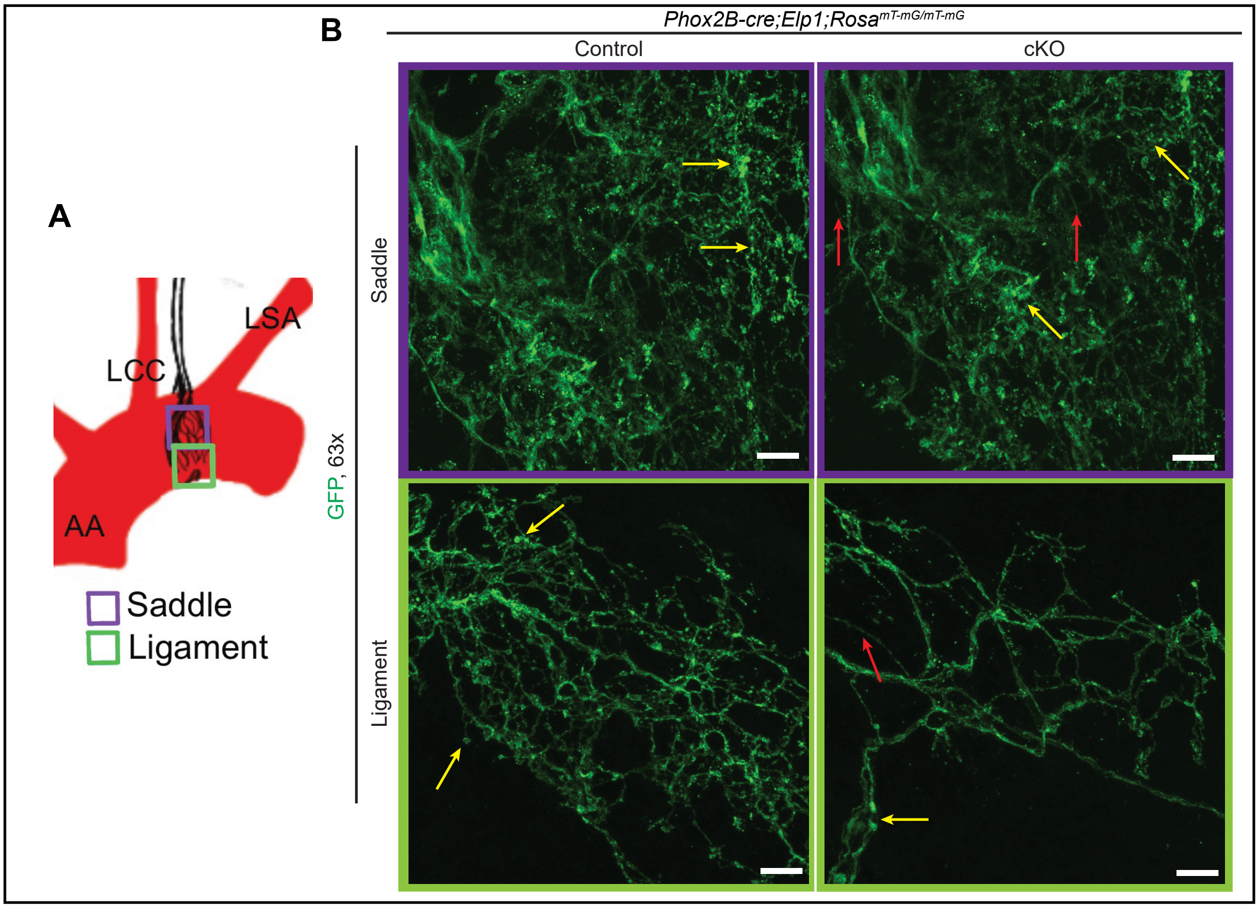


Supplementary Figure 3
